# Impact of deleterious mutations, sexually antagonistic selection and mode of recombination suppression on transitions between male and female heterogamety

**DOI:** 10.1101/480749

**Authors:** Paul A. Saunders, Samuel Neuenschwander, Nicolas Perrin

## Abstract

Deleterious mutations accumulating on non-recombining Y chromosomes can drive XY to XY turnovers, but are thought to prevent XY to ZW turnovers, because the latter require fixation of the ancestral Y. Using individual-based simulations, we explored whether and how a dominant W allele can spread in a young XY system that gradually accumulates deleterious mutations. We also investigated how sexually antagonistic (SA) polymorphism on the ancestral sex chromosomes, and the mechanism controlling X-Y recombination suppression affect these transitions. In contrast with XY to XY turnovers, XY to ZW turnovers cannot be favored by Y chromosome mutation load. If the arrest of X-Y recombination depends on genotypic sex, transitions are strongly hindered by deleterious mutations, and totally suppressed by very small SA cost, because deleterious mutations and female-detrimental SA alleles would have to fix with the Y. If, however, the arrest of X-Y recombination depends on phenotypic sex, X and Y recombine in XY ZW females, allowing for the purge of Y-linked deleterious mutations and loss of the SA polymorphism, causing XY to ZW turnovers to occur at a neutral rate. We generalize our results to other types of turnovers (*e*.*g*., triggered by non-dominant sex-determining mutations) and discuss their empirical relevance.

## Introduction

Several lineages of plants and animals (including birds, mammals, and *Drosophila*) present highly differentiated sex chromosomes, where a large, gene-rich X (or Z) chromosome contrasts with a small, gene-poor and degenerated Y (or W) chromosome. The canonical model of sex chromosome evolution assigns a crucial role to sexually antagonistic (SA) mutations to the first step of the process (Fisher, 1931; Charlesworth and Charlesworth, 1980; Rice, 1984) : in the case of a young XY system, a male-beneficial mutation occurring on the proto-Y chromosome should spread, even if highly detrimental to females, because it is more likely to be transmitted to sons than to daughters. As a second step, recombination will be suppressed between this SA locus and the sex-determining locus (*e*.*g*., through an inversion), generating strict co-segregation of the two loci (Rice, 1987). As a side effect, however, recombination arrest will induce the accumulation of deleterious mutations on the Y chromosome, followed by gene loss and degeneration, and possibly accompanied by dosage compensation (Bergero and Charlesworth, 2009).

However, many other lineages (including in teleost fish, amphibians and non-avian reptiles) present homomorphic sex chromosomes. Two main reasons have been invoked to account for this lack of differentiation. First, if genetic control over sex determination is not complete (allowing for occasional sex-reversal), and recombination arrest depends on phenotypic sex, then X and Y chromosomes will recombine occasionally in sex-reversed XY females, preventing sex-chromosome differentiation in the long term (the ‘fountain-of-youth’ model; Perrin 2009; Rodrigues et al. 2018). Second, sex chromosomes in these lineages might show high rates of turnovers, whereby a new pair replaces the ancestral sex chromosomes before they have had time to degenerate (Schartl, 2004; Volff et al., 2007). Several processes may drive such turnovers. Bull and Charnov (1977) proposed over four decades ago that turnovers could be mediated by genetic drift. So-called neutral turnovers, when driven by sex-determining mutations that are epistatically dominant (*i*.*e*., that override the action of the resident sex-determining gene), are more likely to occur than substitutions at neutral autosomal loci, and turnovers that induce a change in the heterogamety pattern (*e*.*g*., transition from an XY system to a ZW system) tend to become more likely as effective population size (*N*_*e*_) decreases (Veller et al., 2017). Nevertheless, neutral turnovers that maintain the system of heterogamety (*e*.*g*., XY to a new XY system) are 2-4 times more likely to occur than the former (except under extremely low *N*_*e*_; Saunders et al. 2018). One reason is that an XY to XY turnover requires the fixation of the ancestral X chromosome as an autosome, while an XY to ZW turnover requires the fixation of the ancestral Y. The initial difference in frequency between the X and the Y (0.75 *vs*. 0.25), makes the fixation of the X chromosome by chance more likely. A second potential driver for turnovers is SA selection : if a mutant sex determiner arises in close linkage with a SA allele beneficial for the sex determined by that mutation (*e*.*g*., mutant male determiner in close linkage with a male beneficial allele), that newly arisen determiner will have a selective advantage, as the male beneficial - female detrimental allele will be more likely to be inherited by sons than daughters. SA selection was shown to be able to facilitate both transitions that maintain the heterogamety pattern, and transitions that change it (van Doorn and Kirkpatrick, 2007, 2010). A third potential driver of turnovers is the load of deleterious mutations on non-recombining Y (or W) chromosomes. Blaser et al. (2013, 2014) formalized this model for an XY to XY transition, and showed that, given specific combinations for the coefficients of selection (*s*) and dominance (*h*) of deleterious mutations accumulating on the Y, as well as effective population size *N*_*e*_, the benefits of fixing a new, mutation-free Y chromosome can outweigh the cost of losing male-beneficial mutations fixed on the ancestral Y chromosome.

This latter mechanism has been suggested to hinder XY to ZW transitions, because the ancestral Y fixes as an autosome once the transition is over, which is obviously detrimental if it is loaded with deleterious mutations (Blaser et al. 2013, 2014). However, theoretical evidence has been provided only for cases where deleterious mutations have a strong negative effect on fitness, and are completely recessive (van Doorn and Kirkpatrick, 2010; Veller et al., 2017). While this situation is well suited to account for systems in which the degeneration of the Y has advanced to the point where YY individuals bear large fitness costs, it is less applicable to young sex chromosome systems which have had less time to accumulate deleterious mutations. In this paper, we model the evolution of young non-recombining sex chromosomes (XY), which gradually accumulate deleterious mutations and carry a SA locus. We use individual-based simulations to explore quantitatively whether and how different values of *h*, *s* and *N*_*e*_ effectively prevent XY to ZW turnovers.

Additionally, we explore whether and how this outcome depends on the mechanism arresting recombination between the sex chromosomes. This arrest can result from a sex-specific recombination suppression genome-wide, as documented in *Drosophila* (male achiasmy; Morgan 1914) and some Lepidoptera (female achiasmy; Tanaka 1914). Note that recombination is not necessarily entirely suppressed in the heterogametic sex : in some amphibians, including the European tree frogs *Hyla arborea* and the common frog *Rana temporaria*, male recombination is restricted to the tips of chromosomes, and largely absent in their center (Rodrigues et al. 2013; Brelsford et al. 2016; Jeffries et al. 2018). Alternatively, suppression can be specific to the sex chromosome pair : caused by a chromosomal rearrangement (*e*.*g*., an inversion; Kirkpatrick 2010), or a gradual spread of the non-recombining region (*e*.*g*., through a change of the location of cross-over events; Charlesworth et al. 2005). A profound difference between the two modes of recombination suppression (genome wide or chromosome specific) is that in the former, recombination between the two sex chromosomes can occur in sex-reversed XY females (*e*.*g*., Rodrigues et al. 2018), which might affect XY to ZW turnovers as it gives raise to the possibility for X-Y recombination in XY ZW females.

## Methods

For all simulations, we used a genetic model and chromosomal architecture similar to those in Blaser et al. (2013), and identical parameter values, which allowed us to directly compare the effect of Y chromosome mutation load on the dynamics of sex chromosome turnovers that either change the pattern of heterogamety, or preserve it.

### Genetic model

#### Sex determination

The genomic model consists of one pair of sex chromosomes and one pair of autosomes (fig. S1). The sex chromosomes (X and Y) harbor the resident sex-determining locus with two alleles, *X* and *Y*. Sex determination is strictly genetic, so that *XX* individuals are females and *XY* individuals are males. The autosomal pair harbors a gene involved in the sex-determination cascade, initially fixed for an allele *f*. This *f* allele can mutate to a feminizing allele *F* that overrides the masculinizing effect of allele *Y* (epistatic dominance), so that *XY fF* individuals develop as females. If both loci are polymorphic, males can be *XY ff* or *YY ff*, and females *XX ff*, *XX fF*, *XY fF* or *YY fF*. A sex chromosome turnover is achieved once polymorphism at the ancestral sex-determining locus is lost, with all males being *YY ff* and all females *YY fF* (*i*.*e*., ZZ and ZW).

#### Deleterious mutations

Both pairs of chromosomes carry 10 loci evenly spaced (10 cM) around the sex determining loci, which can mutate to a deleterious form. The effect on fitness of a single deleterious mutation depends on a selection coefficient *s* and a dominance coefficient *h*, so that the fitness of an individual heterozygous for that locus is reduced by *hs* and that of an individual homozygous for the mutant allele by *s*. Deleterious mutations affect the mutation load component of fitness (*w*_*M L*_) which is assumed to be :

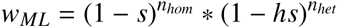

where *n*_*hom*_ and *n*_*het*_ are, respectively, the number of loci homozygous and heterozygous for a deleterious mutation.

#### SA selection

In some of our simulations, the sex chromosome pair carries a bi-allelic sexually antagonistic locus (*e*.*g*., a coloration gene), inserted between the sex-determining locus and one of the two adjacent loci subject to deleterious mutation (genetic distance : 10cM, fig. S1). The two alleles are a male beneficial - female detrimental allele (*a*^*m*^), and a female beneficial - male detrimental allele (*a*^*f*^). The sexually antagonistic component of fitness (*w*_*S A*_), depends on the phenotypic sex and genotype of individuals at that locus : *w*_*S* *A*_ = 1 in *a*^*m*^*a*^*m*^ males and *a* ^*f*^ *a* ^*f*^ females, *w*_*S* *A*_ = 1 - 2*c* in *a* ^*f*^ *a* ^*f*^ males and *a*^*m*^*a*^*m*^ females and *w*_*S* *A*_ = 1 - *c* in all *a*^*f*^ *a*^*m*^ heterozygotes, where *c* is the cost of SA selection (*i*.*e*., the impact on fitness of bearing a detrimental allele). In the presence of this locus, the total fitness of an individual (*W*) is the product of its two components

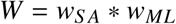

#### Recombination pattern

Recombination depends on either phenotypic sex or genotypic sex. In the first case, recombination occurs in females and never in males, regardless of their genotype. This mimics species in which recombination is either arrested or largely restricted genome-wide in the heterogametic sex (*e*.*g*., frogs, *Drosophila*). In the second case, all loci on the Y chromosome are assumed to be included in an inversion, so that recombination occurs only in individuals homozygous at the ancestral sex-determining locus (*XX* and *YY*), regardless of phenotypic sex. In practice, we implemented a bi-allelic locus controlling recombination rate. It is strictly linked to the *XY* sex-determining locus in all individuals (0 cM, fig. S1), and allows recombination either only in females regardless of genotypic sex, or only in homozygotes regardless of phenotypic sex. The recombination rate (*r*) depends on the genetic distance in cM (*m*) following Haldane’s mapping function (Haldane, 1919) :

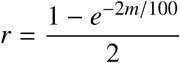

In individuals in which recombination is not allowed, *r* = 0.

### Simulations

Individual-based simulations were run with quantiNemo 2 (Neuenschwander et al., 2018). Unless mentioned otherwise, all simulations were run for 10^5^ generations with effective population size set to *N*_*e*_ = 10^3^. At each generation (non-overlapping), gametes were drawn with a probability proportional to individual fitness *W*, and gametes were paired randomly (one male and one female) to constitute the *N*_*e*_ individuals of the next generation (soft selection). The initial frequency of *X* and *Y* alleles on the sex chromosome pair was set to 0.75 and 0.25 respectively. On the autosomal pair, the sex-determining locus was initially fixed for the *f* allele, and allowed to mutate to its dominant form *F* at a rate *μ* = 10^-5^, with no back mutation possible. A first set of simulations was run to evaluate how the mode of recombination arrest impacts sex chromosome turnovers (in the absence of SA selection : SA locus absent). The loci subject to deleterious mutation were allowed to mutate to their deleterious form at a rate *μ* = 10^-4^. The coefficients of deleterious mutations varied independently between sets of simulations (with values *s* = {0.001, 0.0025, 0.005, 0.01, 0.02, 0.03, 0.04, 0.05, 0.1, 0.15, 0.2} and *h* = {0, 0.1, 0.15, 0.25, 0.5, 0.75, 1}). In a single run, *h* and *s* were the same for all loci. In a similar manner to Blaser et al. (2013), we assessed, for each combination of parameters, the proportion of simulations in which a transition had occurred at the end of the 10^5^ generations (across 100 replicates). We also assessed the mean time to transition, which is the sum of the waiting time until a mutation *F* appears and time for its “fixation” (because the mutation rate was chosen arbitrarily, we were interested in relative time to turnover, rather than its absolute value). In a second set of simulations, we changed the effective population size to *N*_*e*_ = 10^2^ and 10^4^, in order to evaluate the effect of drift on these turnovers. A third set of simulations aimed at determining the impact of a SA gene located on the resident sex chromosomes on the dynamics of XY to ZW transitions in the presence of deleterious mutations. The SA locus was therefore inserted close to the SD locus (fig. S1), with the *a*^*m*^ allele initially fixed on the Y chromosome (frequency : 0.25) and the *a* ^*f*^ allele fixed on the X (0.75). First, we ran simulations with *c* = 0.005, 0.02 across the whole range of *h* x *s* combinations (again, to allow direct comparison with Blaser et al. 2013). As there was no evidence for an interaction between *hs* and *c* values in these simulations, we limited further simulations (aimed at investigating more thoroughly the interaction between SA selection and recombination pattern on turnovers) to a smaller set of parameters : *h* and *s* values were set to 0 (neutral mutations), and *c* varied from 0 to 0.02 with a 0.0025 increment. In case of recombination depending on phenotypic sex, we also explored a larger distance between the SD and SA loci (100 cM, loose linkage; *r* ≈ 0.42). For this set of simulations, the proportion and mean time to transition were assessed, as well as the proportion and mean time to polymorphism loss at the SA locus. Finally, to check the full consistency of our simulations with those from Blaser et al. (2013), we replaced the feminizing allele *F*, by a masculinizing *M* allele and replicated their simulations. Additionally, we also ran simulations with the *M* allele across the whole range of *c* values (with the 0.0025 increment), with either *h* = *s* = 0 (no deleterious mutations), or *h* = 0.15 and *s* = 0.02, so that *hs* = 0.003 (value that maximizes the positive impact of mutation load on XY to XY turnovers; Blaser et al. 2013).

## Results

As expected, the fate of deleterious mutations depends on effective population size as well as dominance and selection coefficients : mutations tend to go to fixation at low *N*_*e*_*hs* values and are eliminated at high *N*_*e*_*hs* values. As also found by Blaser et al. (2013), the shift between these two outcomes occurs between *log*_10_(*N*_*e*_*hs*) = 0 and 1 in the non-recombining region of the Y chromosome (fig. S2). Furthermore, we show that it occurs between *log*_10_(*N*_*e*_ *s*) = 0 and 1 (*i*.*e*., independent of dominance) on the X chromosome (fig. S3). Note that fully recessive mutations (*h* = 0) accumulate on the Y at any *s* and *N*_*e*_ value, being fully sheltered from selection. The two boundaries *log*_10_(*N*_*e*_*hs*) = 1 and *log*_10_(*N*_*e*_ *s*) = 0 (defined above) delimit a window over which mutations accumulate on the Y more than on the X, generating a selective pressure expected to affect the probability of turnovers. The range across which mutations have an effect too strong to accumulate even on the Y (*log*_10_(*N*_*e*_*hs*) > 1) should provide, in the absence of SA selection (*c* = 0), a null expectation for the probability and mean time to sex chromosome turnover, and a reference point to define whether other parameter combinations tend to favor or hinder turnovers. Across this range, turnover probability in our first set of simulations (*N*_*e*_ = 10^3^, *c* = 0) averaged ≈ 0.85 after 10^5^ generations, for any positive *h* value (fig. 1 top panels, right of the dashed vertical line), with a mean time to turnover close to 35,000 generations (fig. S4). This did indeed conform to the neutral expectation (*s* = *h* = 0, red horizontal lines on figs 1 and S4). Fully recessive mutations (*h* = 0), on the other hand, drastically hindered turnovers at any *s* value. Outside of this range (*log*_10_(*N*_*e*_*hs*) < 1), the outcomes of simulations strongly depended on the mechanism of recombination suppression. When recombination was dependent on genotypic sex, on the one hand, turnover probability dropped sharply below *log*_10_(*N*_*e*_*hs*) = 1 (fig. 1 top left panel, left of the dashed vertical line), meaning that the load of deleterious mutations strongly hindered XY to ZW turnovers. The mean time to turnover similarly dropped (fig. S4), showing that XY to ZW turnovers are progressively prevented as deleterious mutations accumulate on the Y (*i*.*e*., turnovers are more likely in young sex chromosome systems, that had less time to accumulate deleterious mutations). Probabilities and time to turnover were higher for small s values, for which mutations also accumulate on the X (*s* = 0.001, *i*.*e*., *N*_*e*_ *s* = 1, regardless of *h*). When recombination was dependent on phenotypic sex, on the other hand, deleterious mutations had no visible effect on the likelihood of turnover (fig. 1, top right panel) and time to turnover (fig. S4), unless mutations were fully recessive (in which case transition probability decreased with increasing values of *s*). The reason the mode of recombination arrest has such a strong impact on turnover probability is that the emergence of a feminizing mutation allows the Y chromosome to be regularly purged through recombination with the X chromosome whenever *XY fF* females appear in the population, but only if recombination depends on phenotypic sex (fig. S5).

**Figure 1:**
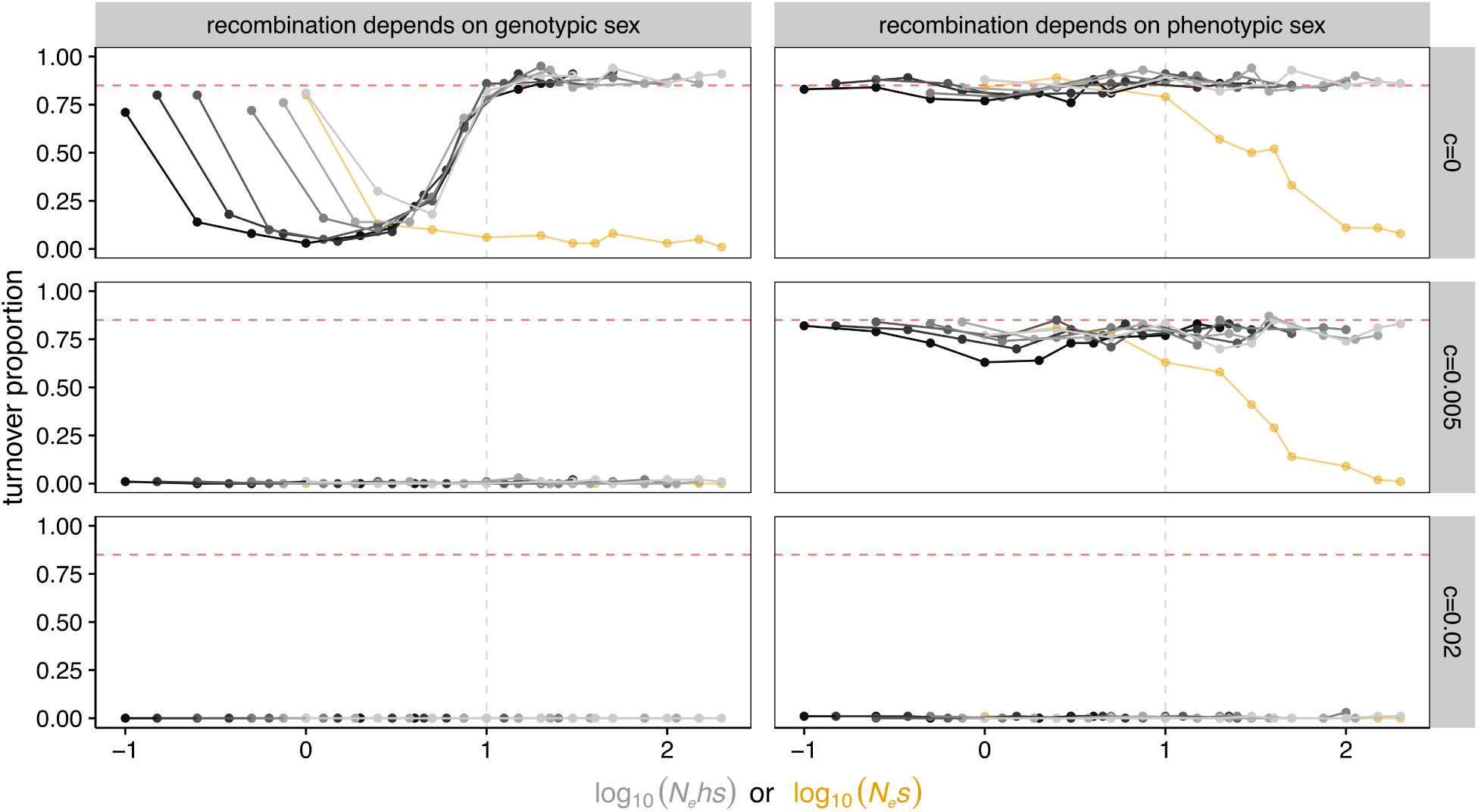
Proportion of replicates (across 100) in which an XY to ZW turnover has occurred at *T* = 10^5^ generations, as a function of *log*_10_(*N*_*e*_*hs*) for *h* from 0.1 to 1 (black to grey scale), or *log*_10_(*N*_*e*_ *s*) for *h* = 0 (yellow), for different strengths of sexually antagonistic selection (*c* = 0, 0.005 and 0.02), and *N*_*e*_ = 1000. The vertical dashed line shows the threshold below which the Y starts accumulating deleterious mutations (*log*_10_(*N*_*e*_*hs*) = 1). The red horizontal dashed line shows turnover proportion (across 100 replicates) for a neutral case (*h* = *s* = *c* = 0).

Varying effective population size had no qualitative effect on turnovers when recombination was dependent on genotypic sex within the parameter spaced tested (figs. S6, S7) : regardless of *N*_*e*_, the likelihood and mean time to turnover dropped at *hs* values that allow deleterious mutations to accumulate on the Y but not on the X (*i*.*e*., between *log*_10_(*N*_*e*_*hs*) = 1 and *log*_10_(*N*_*e*_ *s*) = 0). In contrast, *N*_*e*_ had a qualitative effect when recombination was dependent on phenotypic sex. In populations with large effective size (*N*_*e*_ = 10^4^), even fully recessive mutations (*h* = 0) had no effect on transition likelihood; while in smaller ones (*N*_*e*_ = 10^2^), a drop also occurred at intermediate *hs* values (strongest effect around *log*_10_(*N*_*e*_*hs*) = 0, fig. S6), but with no effect on the mean time to turnover (fig. S7). The reason is that purifying selection is less efficient at keeping deleterious mutations at a low frequency on the X chromosome in populations with smaller effective size (fig. S3), so the Y chromosome cannot be purged as efficiently in sex-reversed females.

The effect of sexually antagonistic selection also varied along with the mode of recombination arrest. When recombination depended on genotypic sex, turnovers were totally prevented by even very weak SA selection (fig. 1, middle and bottom left panels). When recombination depended on phenotypic sex, weak SA selection (*c* =0.005; figs. 1 and S4, middle right panels) had little effect on the likelihood of turnover events, or time until a turnover occurred, compared to simulations without SA selection (*c* = 0) : over all *hs* values > 0, the turnover probability dropped only from 0.85 ± 0.04 (mean ± sd) to 0.78 ± 0.05, though stronger SA selection (*c* = 0.02; fig. 1, lower right panel) completely hindered turnovers.

Broadening the values for *c* showed that, as expected, turnover probability gradually decreases with the strength of SA selection (fig. 2, black lines), and confirmed that the negative effect of SA selection on turnovers is much stronger when recombination suppression depends on genotypic sex rather than phenotypic sex. In the latter case, increasing the genetic distance between the sex-determining locus and the SA locus (from 10 to 100 cM) further increased turnover likelihood and decreased mean time to turnover (figs. 2 and S8), suggesting that recombination between the two loci is what reduces the negative effect of SA selection on transitions from male to female heterogamety.

**Figure 2:**
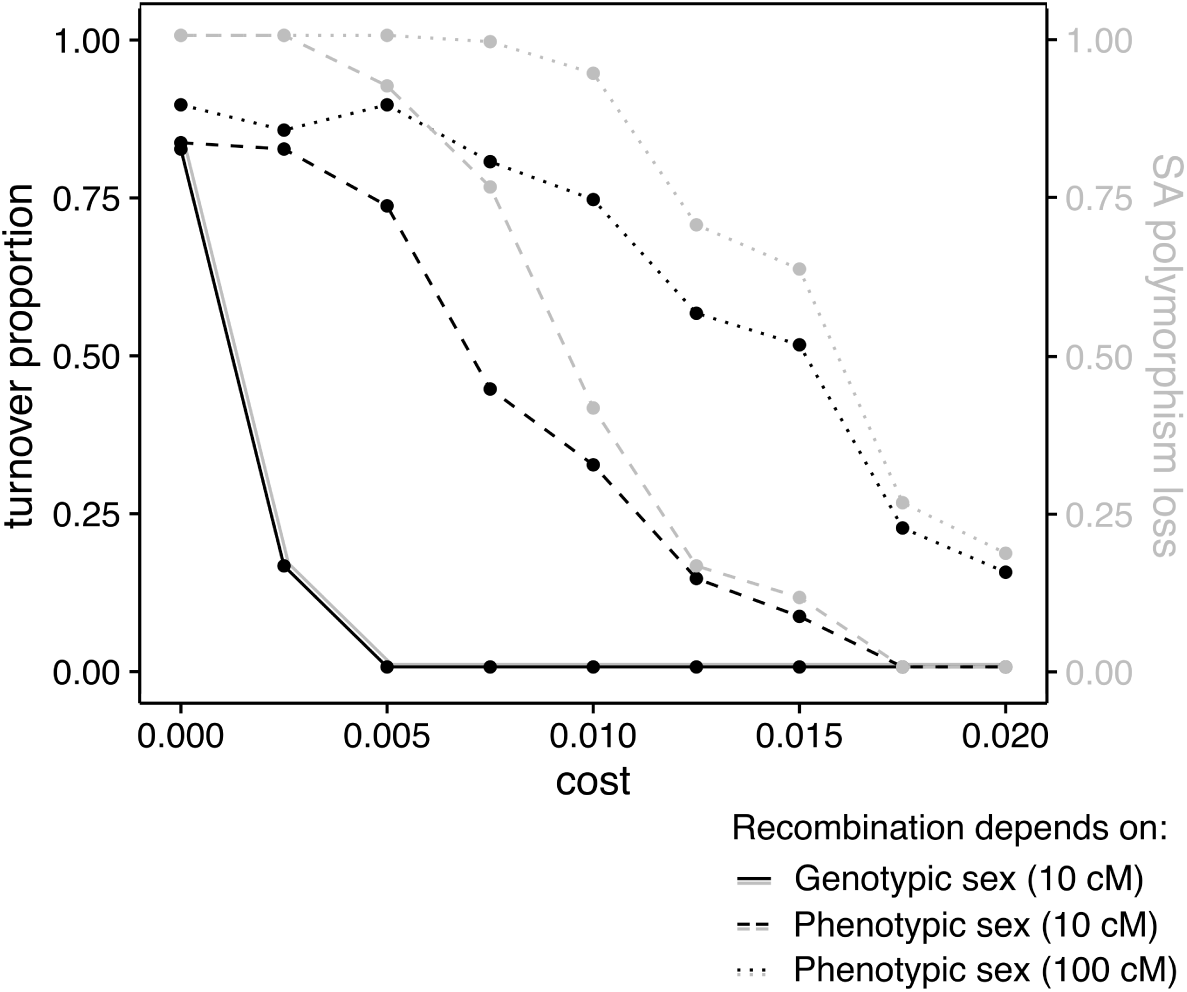
Turnover proportion (black lines) and proportion of simulations during which the polymorphism at the SA locus is lost (grey lines), as a function of the strength of sexually antagonistic selection (cost *c*), across 100 replicates. Parameter values : *N*_*e*_ = 10^3^, *h* = *s* = 0, the values in parentheses in the legend give the distance between the sex-determining locus and the SA locus.

Analyzing the allelic composition of the SA locus at the end of simulations revealed that the probability that one of the two SA alleles goes to fixation also decreases with the strength of SA selection *c* (fig. 2, grey lines) while the mean time to fixation increases (fig. S8). These results are consistent with classical SA theory results showing that SA polymorphism is more easily maintained with stronger SA selection (e.g. Kidwell et al. 1977; Jordan and Charlesworth 2012). Interestingly, mean SA allele fixation time is always shorter than the mean time to turnover, and turnovers in these simulations tend to closely follow the loss of polymorphism at the SA locus (fig. S8, S9). This suggests that turnovers are favored by this loss of polymorphism. Rerunning simulations in which either the male beneficial - female detrimental or female beneficial - male detrimental allele is initially fixed on both the X and the Y chromosomes, brings turnover likelihood back to the neutral expectations (absence of SA locus), regardless of (i) which allele is fixed, and (ii) the value of *c* (fig. S10). This confirms that fixation of either SA allele, made possible by recombination between the sex-determining locus and SA locus in *XY fF* females, is what increases turnover probability. This loss of polymorphism is likely mediated by genetic drift, as supported by the facts that i) loss occurs less frequently at high selection coefficients, and ii) the fixation probability of each of the two alleles is close to their initial frequencies (fig. S11). Thus, although SA variation on the ancestral sex chromosome pair hinders transitions between male and female heterogamety, X-Y recombination in *XY fF* females can break the association between X and Y chromosomes and female- or male-beneficial SA alleles. If either allele is lost by genetic drift, the transition probability increases, and goes back to the expected transition probability in the absence of a SA locus.

To better compare our results to those from Blaser et al. (2013), we re-ran some simulations replacing the autosomal feminizing mutant allele *F* by an epistatically dominant masculinizing allele *M* (note that the mode of recombination arrest cannot impact the spread of the latter, as no XY females are produced throughout XY to XY turnovers). As expected, in the absence of SA variants linked to the resident sex-determining locus, XY to XY turnovers occur with a higher probability than XY to ZW ones, especially if recombination depends on genotypic sex (fig. S12). In the presence of SA selection, this difference is exacerbated for *log*_10_(*N*_*e*_*hs*) values ranging 0 to 1 (fig. S12), and increases with the selective cost of the SA alleles (fig. S13).

## Discussion

As shown by Blaser et al. (2013, 2014), in a male heterogametic sex-determining system, the load of deleterious mutations accumulating on the non-recombining Y chromosome can promote the spread of an emergent male sex-determining mutation, thereby causing a transition in the sex determination mechanism : an XY to XY turnover. In sharp contrast, our present results show that this Y chromosome mutation load cannot favor the spread of an emergent female sex-determining mutation, and therefore does not favor XY to ZW turnovers. Depending on the mechanism that controls recombination arrest, a transition to a ZW system can either be strongly counter-selected, or just allowed to spread at a near neutral rate. Firstly, if recombination arrest depends on genotypic sex (*e*.*g*., caused by a chromosomal inversion on the Y), mutations that fall within the interval *N*_*e*_ *s* = 1 and *N*_*e*_*hs* = 10 strongly hinder turnovers (figs. 1 and S4, top left panels). This corresponds to the interval in which purifying selection is too weak to prevent fixation of deleterious mutations on the Y (fig. S2), but strong enough to prevent their fixation on the X (fig. S3). As thoroughly discussed in Blaser et al. (2013) *N*_*e*_*hs* values that facilitate the accumulation of mutations on the Y (1 < *N*_*e*_*hs* < 10), are likely to be met in nature in many vertebrates species, and it has been confirmed empirically that non-recombining Y chromosomes tend to accumulate deleterious mutations at a faster rate than their X counterpart (*e*.*g*., Berlin and Ellegren 2006; Bergero and Charlesworth 2011; Zhou and Bachtrog 2012), providing the right conditions for Y-deleterious load to have an impact on the spread of sex determiners. Secondly, if recombination arrest depends on phenotypic sex (*e*.*g*., in link with a genome-wide restriction of recombination in males), the load of deleterious mutations on the Y has far less impact on transitions (figs. 1 and S4, top right panels). The reason for this difference is that an XY to ZW turnover requires fixation of the Y chromosome as an autosome, which is costly if it is loaded with deleterious mutations. However, along the process, *XYF f* females are produced, providing an opportunity for X-Y recombination, and therefore purge of the Y deleterious load, but only if recombination depends on phenotypic sex. Our results also suggest that this opportunity for recombination can limit the negative effect that SA selection on resident sex chromosome has on turnovers (fig. 1 middle and bottom panels). Indeed, if recombination depends on genotypic sex, even the weakest cost of bearing a male-beneficial allele on the Y (*c* = 0.005) is enough to greatly limit the spread of female sex-determining mutants. If recombination depends on phenotypic sex, conversely, recombination in *XYF f* females allows decoupling alleles at the sex-determining locus and SA locus, which can lead to the loss of SA variation by genetic drift. Note that our SA model is symmetric (the cost of being homozygous for the detrimental allele is the same in males and females) and that the effect of SA allele is additive. It might seem more plausible that SA alleles are partially dominant in the sex in which they are advantageous, which, according to analytical models, should increases the stability of SA polymorphism (Fry, 2010). While this might decrease the likelihood of XY to ZW turnovers when recombination depends on phenotypic sex by making it harder to fix SA alleles, it does not affect our results qualitatively : SA polymorphism can be lost only in this case, and not if recombination depends on genotypic sex.

The direct comparison of XY to ZW and XY to XY turnover likelihood in our models (fig. S12) suggests that if the sex chromosome deleterious load plays a predominant role in sex chromosome turnovers, transitions that maintain the heterogamety pattern should be more frequent than transitions along which the heterogametic sex switches, especially if recombination suppression depends on genotypic sex (and assuming mutant male and female determiners emerge at the same rate). It is worth noting, also, that in the absence of deleterious mutations, a SA polymorphism on the resident sex chromosomes has the potential to favor XY to ZW transitions over XY to XY ones, provided recombination depends on phenotypic sex (fig. S13). This arises because such a polymorphism, which normally hinders transitions, can be lost through X-Y recombination in *XY Ff* females, generated only in transitions between male and female heterogamety. These observations suggest that it might be possible to assess the relative impact of different evolutionary forces on sex chromosome turnovers (*e*.*g*., SA selection *vs*. deleterious load) by examining the rates of different types of transitions across phylogenies in taxa that experience frequent sex chromosome turnovers. For instance, an unprecedented rate of sex chromosome turnovers has recently been described in Ranidae frogs (Jeffries et al., 2018), and most turnovers have preserved the ancestral mode of male heterogamety, which was interpreted as supporting a role for the load of deleterious mutations (as opposed to SA selection) in driving turnovers. As our present results show, however, this load should not entirely prevent transitions to female heterogamety, given that the lack of XY recombination is controlled by phenotypic sex in Ranidae (Rodrigues et al. 2018). Thus, a role for SA genes cannot be excluded here : as our results also show, turnovers are still much more likely to maintain male heterogamety under the range of deleterious mutations considered (*i*.*e*., between *N*_*e*_ *s* = 1 and *N*_*e*_*hs* = 10) in the presence of male-beneficial alleles on the ancestral Y chromosome (fig. S13). Alternatively, given the genome-wide restriction of recombination in males, an autosomal male-beneficial mutation might be more likely to attract a new sex-determining gene than a female-beneficial mutation, since the new sex determiner will then immediately benefit from full linkage with the SA gene (Sardell et al., 2018). Turnovers towards systems where the sex showing reduced recombination is the heterogametic one might thereby be favored by SA selection, regardless of whether they involve a change in the heterogamety or not. Whether SA selection also plays a role in the sex chromosome turnovers of Ranidae requires further investigation. The rate of characterization of sex chromosome systems has drastically increased in the last few years thanks to the development of genomic technologies, revealing that many taxa experience frequent changes in sex determination mode (Gamble et al., 2015; Blackmon et al., 2017; Gammerdinger et al., 2018; Pennell et al., 2018). The record is nevertheless still too scarce in most groups to quantify the rates of different types of turnovers, but as data keep accumulating, it might become possible in the near future to test more thoroughly our hypotheses.

So far, we have only discussed sex chromosome turnovers driven by epistatically dominant autosomal mutations, occurring in an initially male heterogametic system. Our results can be readily expanded to other types of turnovers. Firstly, the system is symmetric regarding the reverse ZW to XY transition. The same mechanism would allow purging the W from its load of deleterious mutations, provided recombination is arrested in females, not in males (as documented in Lepidoptera). Secondly, the mechanism would also apply to so-called “homologous” turnovers, where the sex-determining mutation arises on the resident sex-chromosome pair, not on an autosome (as documented *e*.*g*., in *Glandirana rugosa*; Miura 2008). A mutation on the X making it a dominant female-determining sex chromosome (W), for instance, would produce WY females in which Y can recombine and be purged before its fixation as a Z chromosome. In contrast, a mutation on the X making it a dominant male-determining allele (a new Y) would induce a homologous XY to XY turnover during which the ancestral Y chromosome is eliminated and no XY females are produced (Bull and Charnov, 1977). Therefore, homologous turnovers that either change or maintain the ancestral pattern of heterogamety should be impacted in a similar manner to their non-homologous counterparts by the accumulation of deleterious mutations and SA selection. Thirdly, transitions may occur through the spread of mutations that are not epistatically dominant over the resident sex determiner (Bull and Charnov 1977). For instance, a transition from male to female heterogamety can be caused by the spread of a “weakly male-determining mutation” *M* in an originally male-heterogametic system (*i*.*e*., from *XX mm*/*XY mm* to *XX mM*/*XX MM*; case 2B in Bull and Charnov, 1977). No XY females are produced throughout this kind of transition, and the resident Y chromosome is the chromosome lost once the transition is complete. Hence, this suggests that a transition from male to female heterogamety can actually be favored by mutation-load selection, but only if driven by a non-dominant male-determining mutation. Conversely, a transition that maintains the pattern of heterogamety can be driven by the spread of a “mildly feminizing determining allele” *F* in an originally male-heterogametic system (i.e., from *XX ff* /*XY ff* to *YY FF*/*YY fF*; cases 3B1 and 3B2 in Bull and Charnov, 1977). Throughout these turnovers, XY females are produced, causing the ancestral Y to be purged from deleterious mutations before it goes to fixation as an autosome. Such an XY to XY turnover would not be favored (but not prevented either) by mutation-load selection. Therefore, the dominance relationship between sex-determining factors is also a parameter to be taken into account when considering the impact on sex chromosome turnovers of deleterious mutations, SA polymorphism, and mode of recombination suppression.

A major finding of our study is that the mode of recombination suppression can have a crucial impact on the spread of sex determining mutations. The ultimate and proximal causes of sex chromosome recombination arrest are still poorly understood (Wright et al., 2016; Ponnikas et al., 2018), and although many sex chromosomes have been shown to carry a non-recombining region, it is still unknown how frequently recombination arrest depends on genotypic or phenotypic sex. Inversions are found on many non-recombining sex-chromosomes, and they have naturally been proposed as a potential cause for suppression, but empirical support remains sparse (Bergero and Charlesworth, 2009), and it has been shown that inversions also tend to fix on sex chromosomes following the arrest (*e*.*g*., *Silene latifolia*, Bergero et al. 2008). Interestingly, sex-reversal experiments in a series of organisms, including houseflies (Inoue et al., 1983), crested newts (Wallace et al., 1997), medaka fish (Matsuda et al., 1999; Kondo et al., 2001) and tilapias (Campos-Ramos et al., 2009), clearly show that the patterns of recombination are essentially controlled by phenotypic sex (review in Perrin 2009). Hence, changes in heterogamety certainly occur despite the accumulating load of deleterious mutations in a large range of organisms. However, this conclusion is likely to only apply at the early stages of sex-chromosome evolution. In the absence of turnovers, lack of recombination can lead over time to more profound changes including evolution of structural differences (*e*.*g*., inversions), accumulation of transposable elements and many recessive deleterious mutations, and evolution of dosage compensation (Beukeboom and Perrin, 2014). All these factors should contribute to prevent X-Y recombination, and therefore, a direct change in heterogamety mediated by the spread of an epistatically dominant W mutation.

## Supporting information

## Acknowledgements

We are thankful to Jérôme Goudet and Elisa Cavoto for stimulating discussions and helpful comments, and Andrew Saunders for proofreading. The computations were performed at the Vital-IT (http://www.vital-it.ch) Center for high-performance computing of the SIB Swiss Institute of Bioinformatics. Funding was provided by the Swiss National Science Foundation (Grant Number 31003A_166323) to Nicolas Perrin.

