## Supplementary material for "Impact of deleterious mutations, sexually antagonistic selection and mode of recombination suppression on transitions between male and female heterogamety"

28/11/2018 - *bioRxiv preprint*

**Fig S13.** Effect of SA selection strength on XY to ZW turnovers *vs.* XY to XY turnovers 12

### Supplementary figures

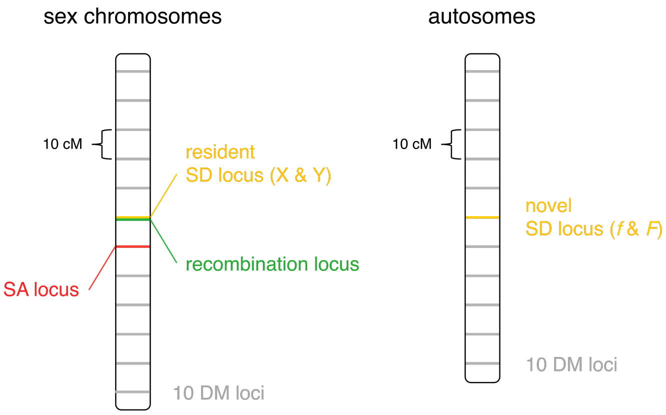

**Figure S1.** Genetic architecture of the two chromosome pairs. SA locus not present in the first two sets of simulations: the genetic distance between the SD locus and closest DM locus underneath it was 10 cM in these simulations. Genetic distances shown are assuming loci are not in a non-recombining region or individual. Abbreviations: SD: sex-determining, SA: sexually antagonistic, DM: deleterious mutation.

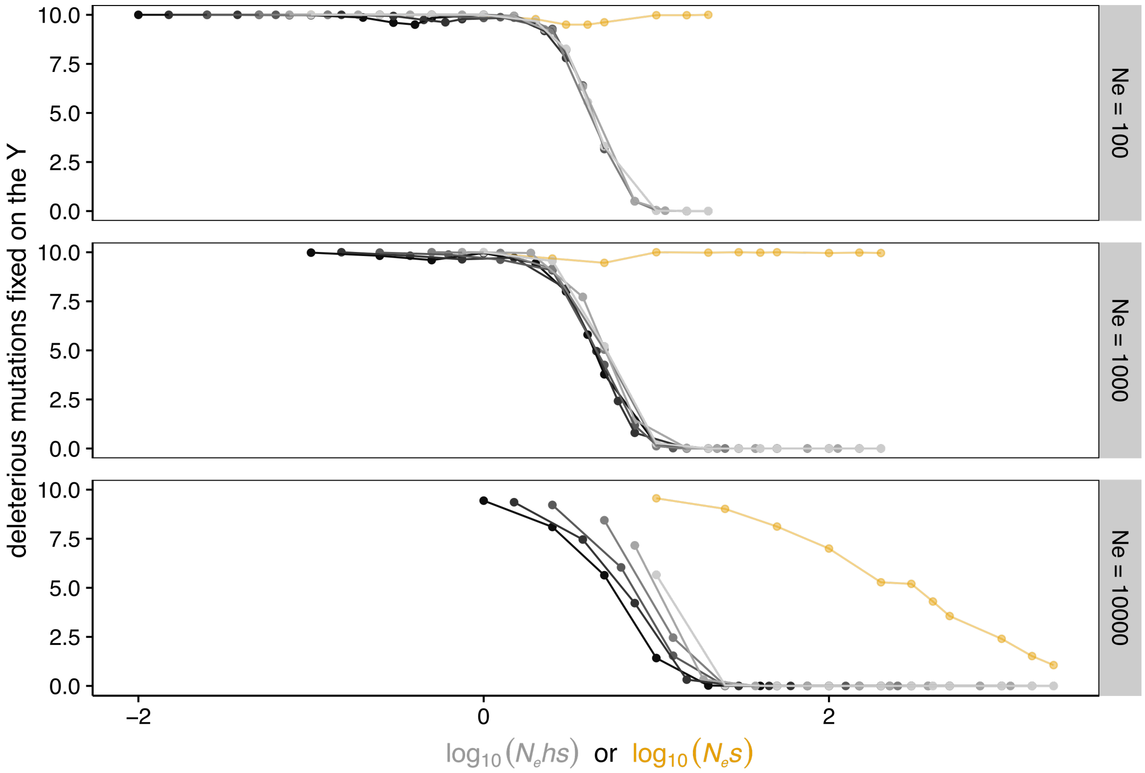

**Figure S2.** Mean number of mutations fixed on the Y chromosome at $T = 10^5$ generations (across 100 replicates), as a function of $log_{10}(N_e hs)$ for $h$ from 0.1 to 1 (black to grey scale), or $log_{10}(N_e s)$ for $h = 0$ (yellow), for different population sizes. In theses simulations, the autosomal $f$ allele was not allowed to mutate to its sex determining $F$ form.

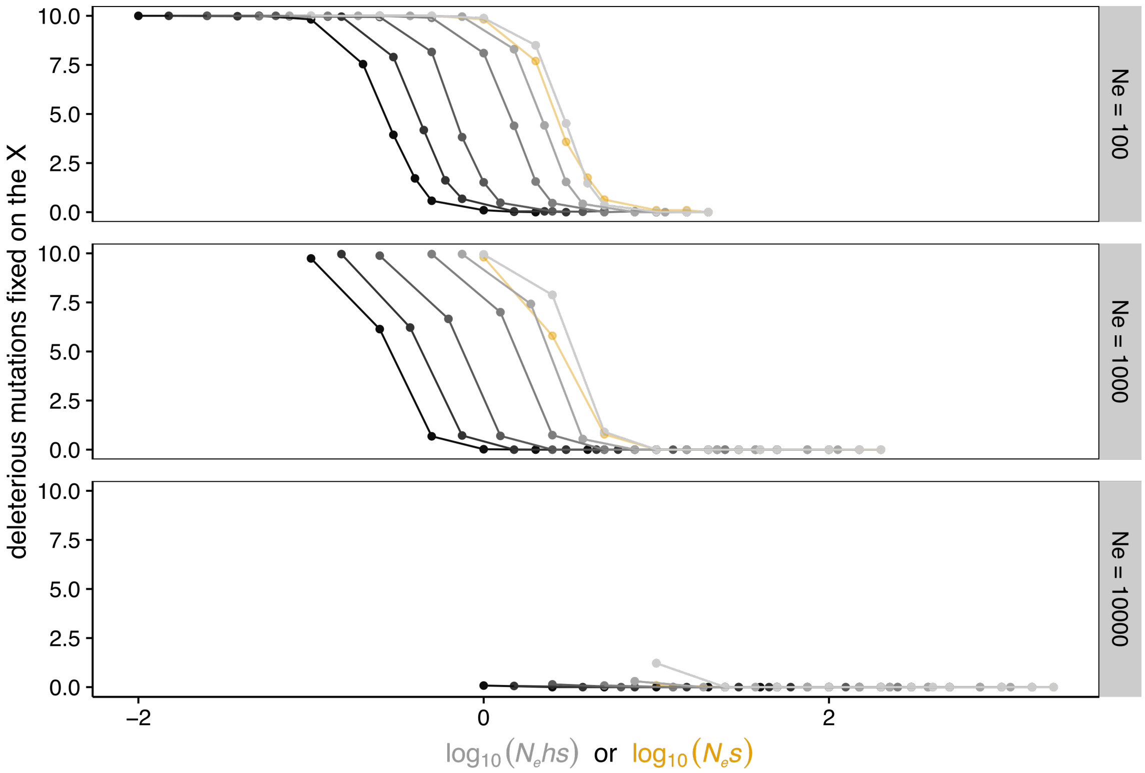

**Figure S3.** Mean number of mutations fixed on the X chromosome at $T = 10^5$ generations (across 100 replicates), as a function of $log_{10}(N_e hs)$ for $h$ from 0.1 to 1 (black to grey scale), or $log_{10}(N_e s)$ for $h = 0$ (yellow), for different population sizes. In theses simulations, the autosomal $f$ allele was not allowed to mutate to its sex determining $F$ form.

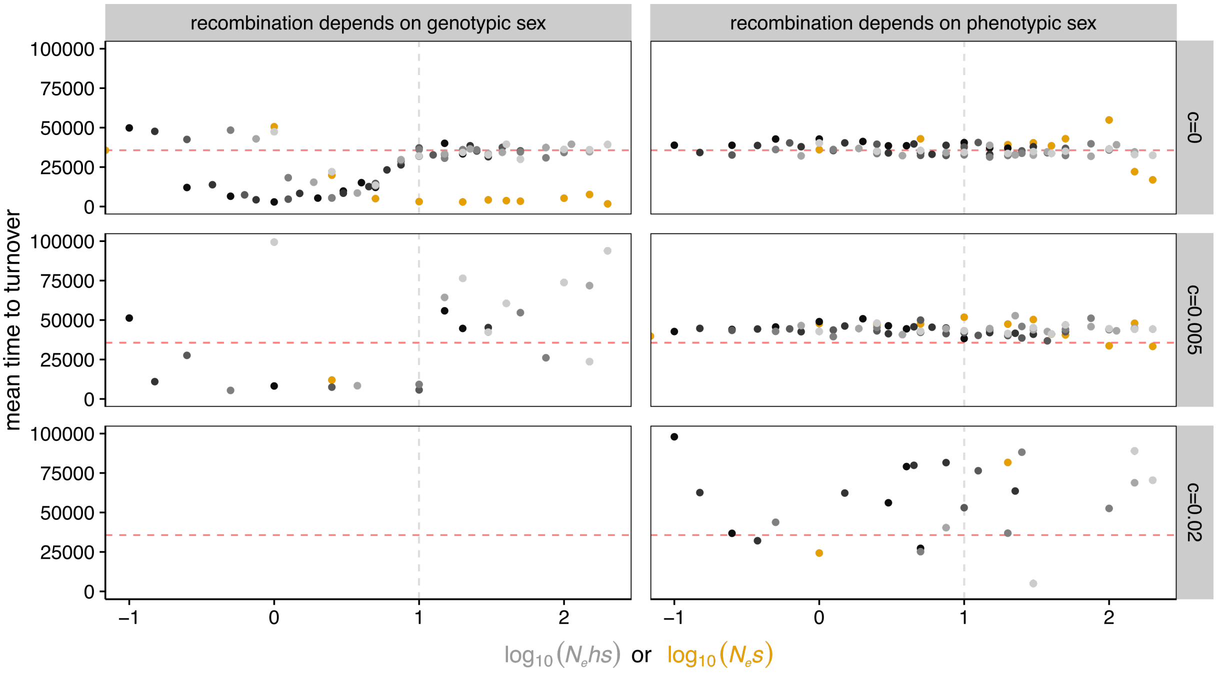

**Figure S4.** Mean time to turnover (across 100 replicates), as a function of $log_{10}(N_e hs)$ for $h$ from 0.1 to 1 (black to grey scale), or $log_{10}(N_e s)$ for $h = 0$ (yellow), for different strengths of sexually antagonistic selection ($c = 0$, 0.005 and 0.02), and $N_e = 1000$. The vertical dashed line shows the threshold below which the Y starts accumulating deleterious mutations ($log_{10}(N_e hs) = 1$). The red horizontal dashed line shows mean time to turnover (across 100 replicates) for a neutral case ($h = s = c = 0$).

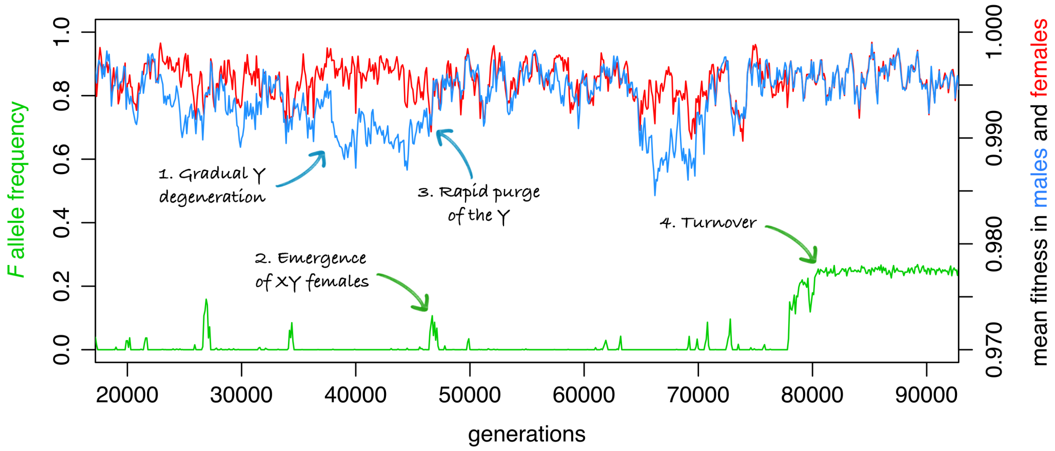

**Figure S5.** Frequency of the feminizing $F$ allele and mean fitness in males and females throughout a part of a single simulation (recombination dependent on phenotypic sex), in which a XY to ZW turnover occurs. Parameter values: $\{N_e = 1000,\ c = 0,\ h = 0.75$ and $s = 0.005\}$.

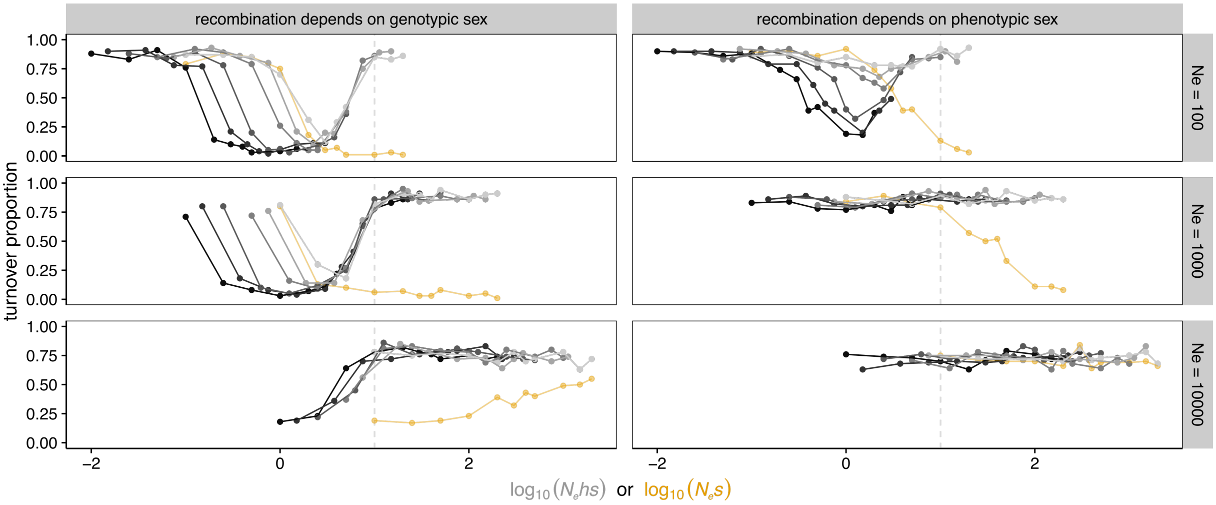

**Figure S6.** Proportion of replicates (across 100) in which a XY to ZW turnover has occurred at $T = 10^5$ generations, as a function of $log_{10}(N_e hs)$ for $h$ from 0.1 to 1 (black to grey scale), or $log_{10}(N_e s)$ for $h = 0$ (yellow), for different population sizes ($c = 0$). The vertical dashed line shows the threshold below which the Y starts accumulating deleterious mutations ($log_{10}(N_e hs) = 1$).

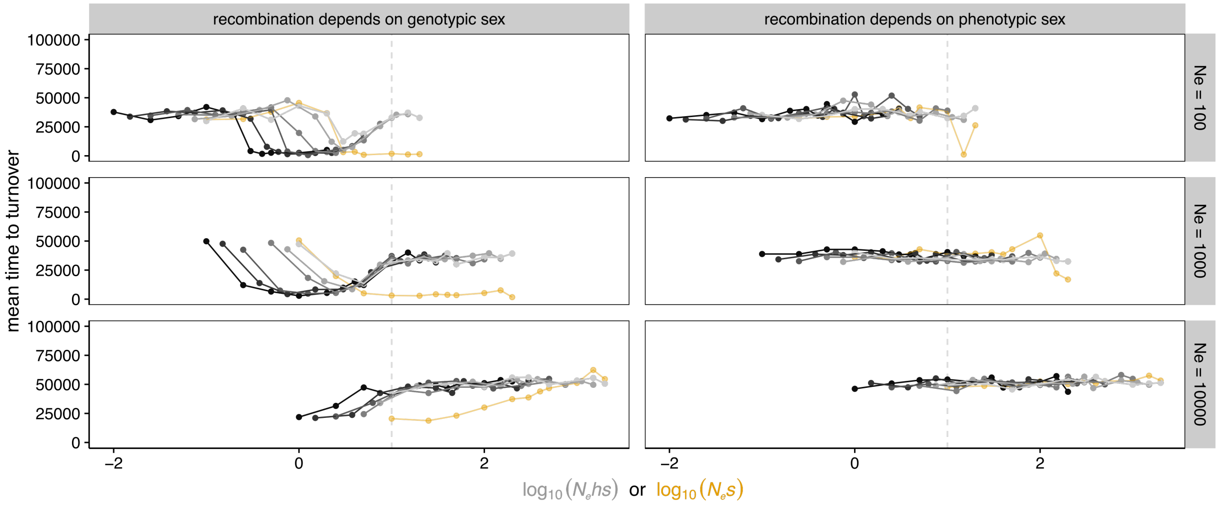

**Figure S7.** Mean time to turnover (across 100 replicates), as a function of $log_{10}(N_e hs)$ for $h$ from 0.1 to 1 (black to grey scale), or $log_{10}(N_e s)$ for $h = 0$ (yellow), for different population sizes ($c = 0$). The vertical dashed line shows the threshold below which the Y starts accumulating deleterious mutations ($log_{10}(N_e hs) = 1$).

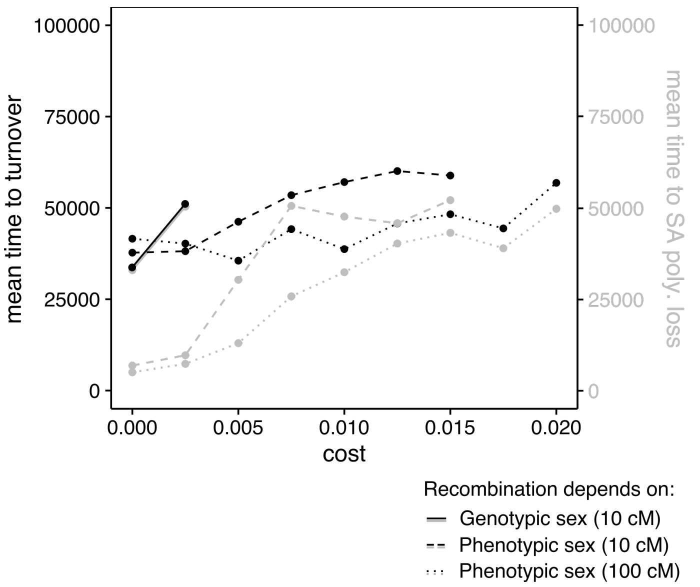

**Figure S8.** Mean time to turnover (black lines) and mean time to loss of polymorphism at the SA locus (grey lines) as a function of strength of sexually antagonistic selection ($c$), across 100 replicates. Parameter values: $N_e = 10^3$, $h = s = 0$, the values in parentheses in the legend are the distance between the sex-determining locus and SA locus.

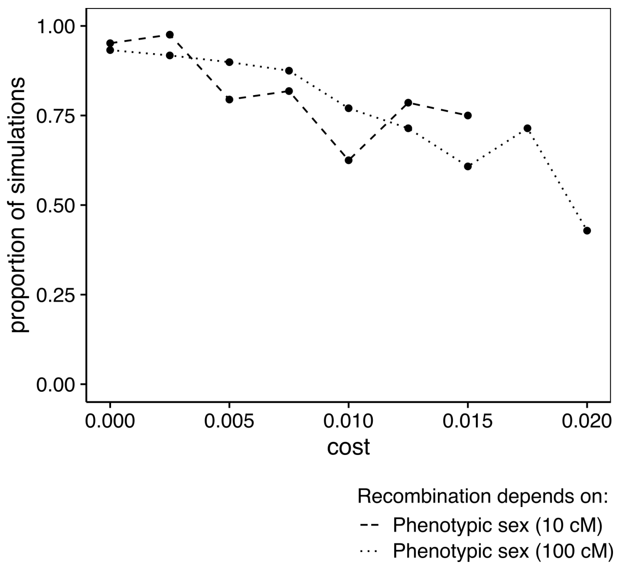

**Figure S9.** Proportion of simulations in which sex chromosome turnover followed loss of polymorphism at the SA locus, across all replicates in which a turnover occurred before the end of the simulation. Parameter values: $N_e = 10^3$, $h = s = 0$, the values in parentheses in the legend is the distance between the sex-determining locus and SA locus.

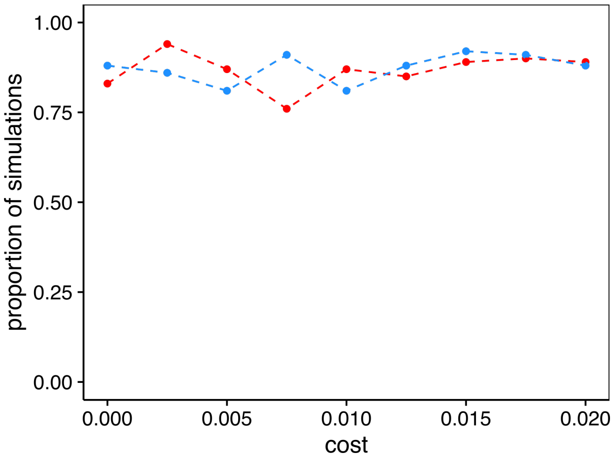

**Figure S10.** Proportion of simulations (across 100) in which a sex chromosome turnover occurred, with one of the two SA alleles initially fixed on both X and Y chromosomes: blue line: $a^m$ fixed, red line: $a^f$ fixed. Parameter values: $N_e = 10^3$, $h = s = 0$, recombination dependent on phenotypic sex, distance between sex determining locus and SA locus: 10cM.

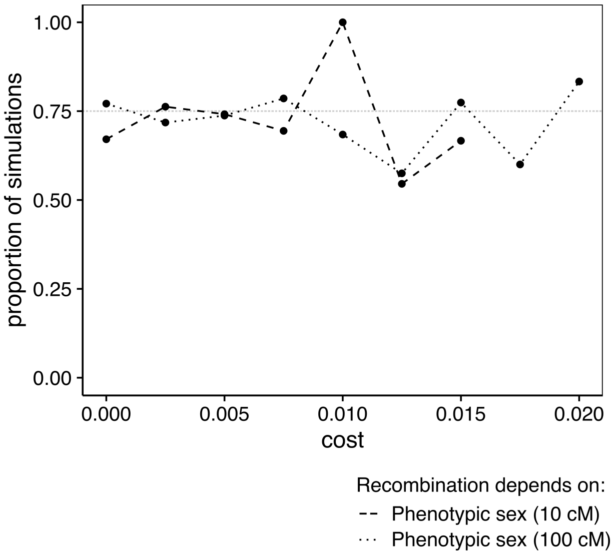

**Figure S11.** Proportion of simulations in which SA allele $a^f$ (initially fixed on the X) went to fixation, across all replicates in which loss of polymorphism at the SA locus preceded sex chromosome turnover. Parameter values: $N_e = 10^3$, $h = s = 0$, the values in parentheses in the legend is the distance between the sex-determining locus and SA locus.

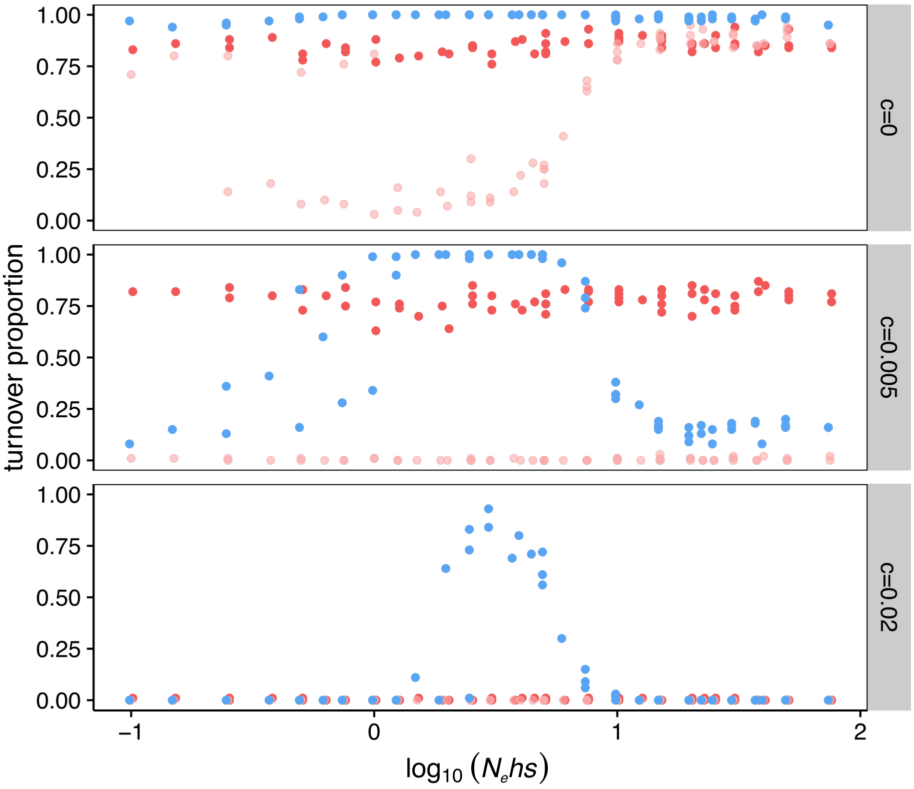

**Figure S12.** Proportion of replicates (across 100) in which a turnover has occurred at $T = 10^5$ generations, as a function of $N_e hs$ (log10 scale, $h$ ranging from 0.1 to 1), for different strength of sexually antagonistic selection ($c = 0$, 0.005 and 0.02), and $N_e = 10^3$. Red dots: XY to ZW turnovers with recombination dependent on phenotypic sex. Pink dots: XY to ZW turnovers with recombination dependent on genotypic sex. Blue dots: XY to XY turnovers.

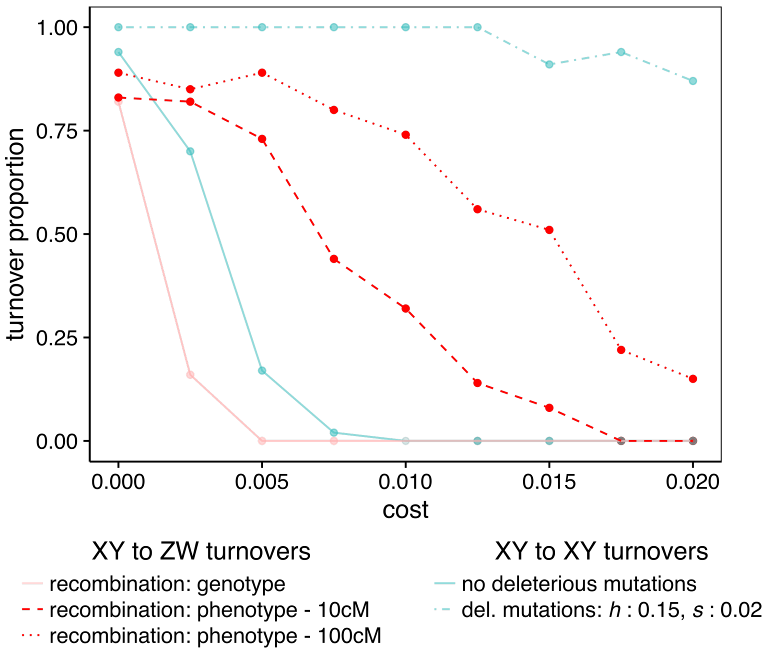

**Figure S13.** Proportion of XY to XY (blue) and XY to ZW (red) turnovers as a function of sexually antagonistic selection strength (cost $c$), across 100 replicates. $N_e = 10^3$. For XY to ZW turnovers, no deleterious mutations accumulated on the sex chromosomes. For XY to XY turnovers with deleterious mutations, values of $h = 0.15$ and $s = 0.02$ were selected because these values maximize the beneficial impact of the accumulation of deleterious mutations on the spread of the masculinizing mutation ($log_{10}(N_e hs) \approx 0.477$).
